# Calcium phosphate nanoclusters modify periodontium remodeling and minimize orthodontic relapse

**DOI:** 10.1101/2024.07.29.605671

**Authors:** Darnell L. Cuylear, Moyu L. Fu, Justin C. Chau, Bhushan Kharbikar, Galateia J. Kazakia, Andrew Jheon, Stefan Habelitz, Sunil D. Kapila, Tejal A. Desai

**Affiliations:** Graduate Program in Oral and Craniofacial Sciences, School of Dentistry, University of California, San Francisco (UCSF), San Francisco, CA, United States; Department of Bioengineering and Therapeutic Sciences, University of California, San Francisco (UCSF), San Francisco, CA, United States; School of Dentistry, University of California, San Francisco (UCSF), San Francisco, CA, United States; Diabetes Center, University of California, San Francisco, San Francisco, CA, United States; Department of Radiology and Biomedical Imaging, University of California, San Francisco (UCSF), San Francisco, CA, United States; Department of Orthodontics and Dentofacial Orthopedics, University of Pittsburgh, Pittsburgh, PA, United States; Department of Preventative and Restorative Dental Sciences, School of Dentistry, University of California, San Francisco (UCSF), CA, United States; Section of Orthodontics, School of Dentistry, University of California, Los Angeles (UCLA), Los Angeles, CA, USA; Department of Bioengineering, University of California, Berkeley (UC Berkeley), Berkeley, CA, United States; School of Engineering, Brown University, Providence, RI, United States

**Keywords:** Orthodontic Relapse, Periodontal ligament, Calcium phosphate, Collagen, Fibrillogenesis

## Abstract

Orthodontic relapse is one of the most prevalent concerns of orthodontic therapy. Relapse results in patients’ teeth reverting towards their pretreatment positions, which increases the susceptibility to functional problems, dental disease, and substantially increases the financial burden for retreatment. This phenomenon is thought to be induced by rapid remodeling of the periodontal ligament (PDL) in the early stages and poor bone quality in the later stages. Current therapies, including fixed or removable retainers and fiberotomies, have limitations with patient compliance and invasiveness. Approaches using biocompatible biomaterials, such as calcium phosphate polymer-induced liquid precursors (PILP), is an ideal translational approach for minimizing orthodontic relapse. Here, post-orthodontic relapse is reduced after a single injection of high concentration PILP (HC-PILP) nanoclusters by altering PDL remodeling in the early stage of relapse and improving trabecular bone quality in the later phase. HC-PILP nanoclusters are achieved by using high molecular weight poly aspartic acid (PASP, 14 kDa) and poly acrylic acid (PAA, 450 kDa), which resulted in a stable solution of high calcium and phosphate concentrations without premature precipitation. *In vitro* results show that HC-PILP nanoclusters prevented collagen type-I mineralization, which is essential for the tooth-periodontal ligament (PDL)-bone interphase. *In vivo* experiments show that the PILP nanoclusters minimize relapse and improve the trabecular bone quality in the late stages of relapse. Interestingly, PILP nanoclusters also altered the remodeling of the PDL collagen during the early stages of relapse. Further *in vitro* experiments showed that PILP nanoclusters alter the fibrillogenesis of collagen type-I by impacting the protein secondary structure. These findings propose a novel approach for treating orthodontic relapse and provide additional insight into the PILP nanocluster’s structure and properties on collagenous structure repair.

## Introduction

In the U.S., approximately 4.8 million people annually receive orthodontic treatment to correct malpositioned teeth^1^. By doing so, orthodontic treatment improves orofacial aesthetics, oral functions such as mastication and speech, as well as decreases the risk of dental decay and gum disease^2–5^. However, a critical limitation impacting treatment outcomes is relapse, or the tendency of teeth to move back to their original position after treatment. Currently, mechanical approaches such as retainers are the primary method to stabilize teeth after orthodontic treatment and rely heavily on patient compliance. More invasive approaches, including fiberotomies and interdental reductions, are less common and have weaker evidence to reduce relapse^6,7^. Nevertheless, relapse occurs in upwards of 67% of patients, likely due to non-compliance with retainers such that the biological processes involving the periodontal ligament (PDL) remodeling and bone turnover reverse the gains of therapy^6,8,9^. Given the critical role both the PDL and bone play in orthodontic relapse, therapeutic approaches that impact one or both tissues would be an ideal method to stabilize teeth after orthodontic therapy.

Orthodontic tooth movement (OTM) proceeds through a process of pressure and tension. Upon application of forces, the tooth shifts within the PDL resulting in PDL compression in some areas and PDL stretching in others^10^. Orthodontic relapse has been thought to be induced by similar biological reactions reoccurring in the opposite direction of the original treatment^11^. Recent studies examining orthodontic relapse biology have delineated important temporal processes that facilitate the progression of orthodontic relapse. During the initial tooth movement, collagen degradation occurred in regions of compression, and it rapidly recovered within 5 days after the removal of the appliances contributing to relapse. This phase of relapse could be modified with hormones and growth factors^12,13^. In the later stages of relapse, the underlying bone quality has been implicated. After tooth movement, the underlying bone density and quality decreases and gradually improves over the course of relapse^14,15^. Additional drug therapies have been shown to improve the bone in earlier phases of relapse^15–20^. Although various drugs have been experimentally implemented for relapse, no therapies have been adopted for use^21^. Therefore, strategies to improve relapse with more biocompatible approaches are necessary to mitigate relapse and facilitate the translation of these therapies.

Polymer-induced liquid precursor (PILP) solutions utilizing polyanionic polymers like poly aspartic acid (PASP) and poly acrylic acid (PAA) have long been recognized for their potential in biologically relevant context^22,23^. First introduced by Oltsza et al., PILP was characterized for its ability to facilitate intrafibrillar mineralization of collagen type-I^24^. The traditional understanding of PILP involves the chelation of calcium and phosphate ions by these polyanionic polymers, preventing premature apatite precipitation and stabilizing liquid-like hydrated amorphous calcium phosphate phase^25,26^.

Recent developments in PILP research have expanded its potential beyond mineralization-focused applications. While early studies primarily focused on achieving optimal concentrations of calcium and phosphate for mineralization purposes, more recent investigations have unveiled broader biological implications, including stimulating bone regeneration in osteoporotic bone or large calvarial defects and osseointegration of titanium implants^27–29^. Notably, this was achieved by combining PASP and PAA at high concentrations to form PILP nanoclusters with elevated calcium and phosphate concentrations. Despite advancements in achieving high-concentration PILPs (HC-PILP), further research is needed to optimize the HC-PILP formulation, considering the intricate interplay between biological and mineralization properties. For soft tissue applications, such as the non-mineralized PDL, minimizing ectopic mineralization while still achieving positive aspects of HC-PILPs is ideal^30,31^. Therefore, this study aims to contribute to this ongoing exploration by investigating the effects of HC-PILP formulations on periodontal tissues and their influence on orthodontic relapse.

## Methods

### Synthesis of High Concentration PILP nanoclusters (HC-PILP)

High concentration PILP nanoclusters (hereafter referred to as HC-PILP) were synthesized following Yao et al. with some modifications^27^. First, 0.1 M solutions of CaCl_2_ and Na_2_PO_4_ were separately prepared in deionized water. Second, 0.3 g/ml PASP (Molecular weight, MW= 14 kDa) was prepared in 1 mL of deionized water and 0.24 g PAA (MW= 450 kDa) was prepared in 0.8 mL water and 4 mL of 0.1 M Na_2_PO_4._ Both the PASP and PAA/Na_2_PO_4_ solutions were sonicated for 10 minutes on ice and then subsequently placed on a rotator at 15 RPM overnight at 37 C. To form the HC-PILP, 2.0 mL of the 1.0 M CaCl_2_ solution was mixed with 0.2 mL of the 0.3 g/mL PASP solution and then 2.4 mL of the PAA/Na_2_PO_4_. Na_2_PO_4_ was added dropwise with rapid mixing at 1200 RPM. This solution was then pH adjusted to 7.4 using 0.3 and 0.1 M NaOH. Samples were subjected to filtration through a 40 μm filter (Steriflip Vacuum Tube Top Filter, Sigma Aldrich, St. Louis, MO). As a control throughout the study, the above protocol was repeated and the CaCl_2_ and Na_2_PO_4_ solutions were replaced with water to produce PAA and PASP only in water (PAA/PASP only). The final concentrations of PASP and PAA were 39.14 and 13.01 mg/ml, respectively. For HC-PILP, calcium and phosphate final concentrations were 41.67 mM. All experiments were performed with this final solution unless otherwise indicated.

### Characterization of HC-PILP Structure, Size, and Biological Responses

To determine the size of the HC-PILP, dynamic light scattering (DLS) and Cryo-electron microscopy (cryo-EM) were performed. After synthesis, HC-PILP were immediately placed in DLS cuvettes (Sigma Aldrich, St. Louis, MO, USA) and measurements were obtained on a Malvern Zetasizer Nano ZS. Particle hydrodynamic radius was obtained using Z_avg_ measurements and reported as number percentage. Cryo-EM micrographs were captured on a Glacios cryo-TEM. Briefly, Quantifoil R1.2/1.3 grids (Quantifoil, Großlöbichau, Germany) were treated by glow discharge. 5 μL of synthesized HC-PILP were added to the glow-discharged grids, blotted for 1 second, relaxed, and vitrified by plunging into liquid ethane at liquid nitrogen temperature (−196 °C). A Fast Fourier Transform (FFT) algorithm was utilized to assess the crystallinity of the sample. To determine the phase composition and functional groups in the samples, x-ray diffraction (XRD) and attenuated total reflectance Fourier transform infrared spectroscopy (ATR-FTIR) were used to analyze the samples, respectively. Prior to analysis, samples were pre-frozen at -80 C for 2 hours, lyophilized for 24 hours, and pulverized with a mortar and pestle at the time of analysis. Diffraction data was obtained on a Rigaku Miniflex 6G Benchtop Powder XRD. The X-ray generator was set at 40 kV and 15 mA with a scan speed of 5.00°/min from 10-70°. ATR-FTIR (Perkin Elmer Spotlight 400) was performed with a scan resolution of 4 cm^-1^ from 4500 to 450 cm^-1^ and a constant anvil pressure value of 45. Using the Spectrum software (Perkin Elmer), a background scan was recorded immediately after each sample scan to facilitate background correction. After background corrections, an ATR correction algorithm was applied.

To understand the deformation dynamics of HC-PILP, rheological measurements were conducted on Rheometer AR2000 (TA Instruments, New Castle, DE, USA). Briefly, to measure the rheological properties, 25 mm parallel plate steel geometry with a gap of 750 μm was used at 37 °C. 370 μl of PILP nanoclusters were dispensed on the base plate, strain sweep and frequency sweep tests were performed. For strain sweep steps, the strain was varied from 0.1% to 1000% at a constant angular frequency of ω =10 rad/s, while for frequency sweep steps, the frequency was varied from 0.1 to 628 rad/s at a constant strain of 10% in the linear viscoelastic region (LVR). Data analysis was performed in Rheology Advantage (TA Instruments, New Castle, DE, USA) to profile the critical rheological parameters including storage modulus (G’), loss modulus (G”), and complex viscosity (|η*|), and Tan (δ).

To examine the extent of mineralization induced by HC-PILP, *in vitro* assays were performed using pre-fibrillated collagen type-I (Advanced Biomatrix, Carlsbad, CA, USA) and MC3T3-E1 (subclone 4) cells (ATCC, Manassas, VA). 1 mg of pre-fibrillated collagen was added to a centrifuge tube with either 1 mL of water, PAA/PASP, or HC-PILP. Samples were incubated at 37 C at 12 RPM for either 7 or 14 days (7D and 14D). After incubation, samples were washed thrice in water, flash-frozen in liquid nitrogen, and lyophilized for 48 hours. Scanning and transmission electron micrographs (SEM and TEM) were obtained to visualize collagen periodicity and mineral formation. For SEM, samples were sputter-coated with iridium to a final thickness of 3 nm. SEM images were done using a 5 kV beam on the Phenom Pharos SEM. For TEM, Formvar/Hexagonal 100 mesh grids (Electron Microscopy Sciences, Hatfield, PA, USA) were glow-discharged and samples were briefly wetted in water and added to the grid to dry overnight at room temperature. Finally, samples were analyzed with ATR-FTIR. As described above, spectra were collected for each sample and then were background- and ATR-corrected using the spectrum software.

MC3T3-E1 mineralization was assessed by Alizarin Red S staining and quantification. MC3T3-E1 (subclone 4) were cultured in *α*-MEM without Ascorbic Acid (AA) supplemented with 10% fetal bovine serum and 1% penicillin/streptomycin. Cells were cultured for at least 2 passages prior to use. Cells were then seeded in a 24-well plate at 5×10^5^ cells/well with *α*-MEM supplemented with 50 µg/mL AA to induce matrix deposition^32–34^. Cells were then treated with 30 µL of PAA/PASP or HC-PILP for 14 or 24 days and media was replaced every 3-4 days. Briefly, after incubation with experimental conditions, the cells were washed with PBS and fixed in ice-cold 4% PFA. Cells were then stained with 40 mM Alizarin red S (pH 4.1) (Sigma Aldrich, St. Louis, MO, USA) for 20 minutes and then washed with fresh deionized water 5 times for 5 minutes. After washing the samples were dried overnight and scanned at 1,200 dpi (Epson perfection V39, Long Beach, CA) and image processed using ImageJ (National Institutes of Health, Bethesda, MD, USA). For semi-quantitative measurements, Alizarin red was extracted by incubating the stained cells in 10% (v/v) acetic acid for 20 minutes. Cells in acetic acid were added to 1.5 mL centrifuge tubes and mineral oil was placed on top of the slurry prior to heating to prevent evaporation during the heating step. Samples were then heated at 85 C for 10 minutes and immediately cooled on ice for 5 minutes. The cooled samples were centrifuged at 20,000g for 15 minutes and 50 μL of each sample was added to a 96-well plate and absorbance was measured at 405 nm. Alizarin red S quantifications were calculated using an established standard curve with known Alizarin red S concentrations.

To investigate the effect of PAA/PASP and HC-PILP on osteoblast differentiation, MC3T3-E1 (subclone 4) cells were used. Cells were seeded into 24-well plates and treated with the same conditions as above. Genetic material was harvested and purified using the RNeasy Mini Kit (Qiagen, Hilden, Germany). Genomic DNA was removed with additional steps using the RNase-Free DNase Set (Qiagen). RNA was converted into cDNA using the iScript cDNA synthesis kit (Bio-Rad Laboratories, Hercules, CA), and a Viia7 qPCR machine (Life Technologies, Carlsbad, CA) was used to measure the relative expression level of the gene targets to the housekeeping gene beta-actin. Expression levels of all genes were evaluated using the Fast SYBR Green Mastermix (Life Technologies, Grand Island, NY) and custom DNA primers (Integrated, Coralville, IA) in triplicate for three biological replicates.

### Rat orthodontic tooth movement model and relapse

The animal studies were conducted after approval from the Animal Care and Use Committee (IACUC), and all the procedures adhered to authorized guidelines and regulations. Studies were conducted on Sprague Dawley rats obtained from Charles River Laboratories (CD rat; Wilmington, MA, USA). Prior to orthodontic spring placement, animals were acclimated to a soft food diet for 7 days. Briefly, adult (12-14 weeks) male rats were anesthetized via inhalant isoflurane and a 60g force 12 mm Nickel Titanium closed-coiled spring (American Orthodontics, Sheboygan, WI, USA) was fastened unilaterally from the maxillary first molar to the ipsilateral incisor with 0.010” stainless steel ligature ties for 28 days^16,17^. To ensure ligature placement, retention grooves were notched into the mesial of the left maxillary molar circumferentially and the gingival margin of the incisor. Ligatures were further secured by bonding to the teeth with flowable composite (L-Pop Self-Etch Adhesive and Transbond Supreme LV Low Viscosity Light Cure Adhesive; 3M/ESPE, St. Paul, MN, USA). After placement of the springs and bonding agents, animals were provided with post-operative analgesics (buprenorphine sustained release). To prevent breakage, lower incisors were trimmed weekly, and springs were readjusted as needed to ensure a horizontal force vector. Animals were weighed and checked daily to ensure appliances were in place during the tooth movement phase and appliances were immediately replaced if broken.

After 28 days of OTM, a single injection of selected experimental agents was administered into the distopalatal surface of the maxillary first molar with a precise custom-microliter 34-gauge syringe (Hamilton Company, Reno, NV, USA). Experimental groups were injected in 30 μL total volume as follows: HC-PILP, PAA/PASP in water (ie., no ion species), or PBS, each with 15-17 rats. After administration of the experimental agents, orthodontic springs and ligatures were immediately removed and relapse was allowed for 24 days. To measure the tooth movement, polyvinyl siloxane impressions were taken of the maxillary teeth to fabricate stone tooth models. The occlusal surfaces of the stone models together with a 100 mm ruler were scanned (Epson perfection V39, Long Beach, CA) at 1,200 dpi and measurements were taken in ImageJ. Tooth movement was measured prior to appliance placement, at appliance removal, and at the day of sacrifice (relapse day 4, 8, 12 [R4, R8, and R12], each timepoint with 5-7 rats per group).

### Micro-computed tomography (μCT)

Orthodontically treated hemi-maxillas were harvested at R4, R12, and R24 to assess the extent of bone quality and quantity with μCT. At the time of collection, samples were fixed in 10% formalin for 48 hours with constant agitation, washed with PBS, and stored at -80 °C in 70% EtOH until analysis. Samples were placed in a 34 mm diameter specimen holder and scanned using a μCT system (μCT50 Scanco Medical, Bassersdorf, Switzerland). Samples were scanned with a voxel size of 10 μm, 70 kVp, 114 μA, 8W, 0.5 mm AL filter, and integration time 500 ms. Analysis was performed using the manufacturer’s evaluation system, and a fixed global threshold of 22% (220 on a grayscale of 0-1000) was applied for maxillary interradicular bone. The region of interest was 1.4 mm (140 slices) beginning at the separation of the roots under the furcation. μCT reconstruction and quantitative analyses were performed to obtain the following structural parameters: bone volume fraction (bone volume (bv)/total volume (tv) as %), trabecular spacing (mm), trabecular number (1/mm^3^), and bone mineral density (mg HA/cm^3^). Additional rats were used as a baseline control for no tooth movement (n=7) and after 28 days of OTM (n=5) for post-OTM (Post-OTM) controls.

### Histological preparation and staining

After μCT, samples were decalcified in 0.5 M EDTA (pH 7.4) for ∼10 weeks at 4°C with constant agitation. Hemi-maxillas were processed for paraffin embedding, and serial sections were cut in the axial plane at 5 μm (2 sections per slide) until a depth of 800 μm and collected on Superfrost Plus Microscope slides. Picrosirius red and Masson trichrome staining were performed to assess the extent of collagen organization, abundance, and composition. Briefly, samples adhered on slides were heated in an oven at 55 °C for 20 min., soaked in xylene for a total of 9 min., rehydrated using an ethanol series (100% for 3 times for 3 min., 95% 1 time for 3 min., and 80% one time for 3 min.), and then soaked in deionized water for 5 minutes. For Sirius red staining, sections were incubated in 0.1% (w/v) Sirius Red (Sigma-Aldrich, St. Louis, MO, USA) in saturated picric acid solution. Next, sections were dipped in 0.5% (v/v) acetic acid water solution, 100% alcohol, and then xylene prior to mounting. For Masson Trichrome, the protocol was followed per manufacturers protocol (Sigma-Aldrich, St. Louis, MO, USA). Hematoxylin and Eosin staining was performed to examine the cellularity and extracellular matrix organization. First, slides were stained in Harris’s Hematoxylin (VWR, Radnor, PA, USA) for 5 min., followed by dipping in deionized water for 1 min., and then dipping in Differentiating solution (Sigma-Aldrich) for 1s for a total of three times. Next, slides were immersed in bluing solution (0.1% sodium bicarbonate, pH ∼8.0) for 30 s followed by dipping in water for 1 min. Slides were then soaked in 70% EtOH for 1 min. and then in Eosin Y (VWR) for 45 s. Sections were then dehydrated using an EtOH series (85%, 95% 100%) for 1 min. each before soaking in xylene and mounting.

### Imaging and quantification

All images were taken using a Nikon 6D optical microscope (NIKON Instruments, Inc., Melville, NY) using 4-40x objectives. Subsequent quantifications were performed using a custom macro script in the Nikon NIS-element imaging software.

For collagen analysis, sections from the ∼2/3 of the root (∼800 microns in depth) of each maxilla were selected from the axial plane of the maxilla and stained with Sirius red to assess the quantity and density of collagen in the PDL space. These sections were imaged under cross-polarized light to visualize the collagen fibers under 20x objective. Representative images for polarized shading corrections were taken prior to the start of each imaging session. The PDL space in the mesial part of the distal buccal root and the distal part of the mesial root were observed to represent the compression and tension sides in relapse, respectively. Intensity measurements [Lower/Upper threshold: 5-10/256] in the region of interest were determined using Nikon NIS-element imaging software.

### Fibroblast gene expression

To investigate the effect of PAA/PASP and HC-PILP on fibroblast gene expression, NIH 3T3 fibroblasts were used. NIH 3T3 mouse fibroblasts (ATCC) were cultured in Dulbecco’s modified Eagle’s medium with 10% fetal calf serum and 1% penicillin/streptomycin. Cells were seeded into 24-well plates (10,000 cells/cm^2^) and allowed to adhere overnight. Media was replaced and then treated with PAA/PASP or HC-PILP. Additional independent experiments were performed with TGF-β1 (10 ng/mL) (PeproTech, Waltham, MA, USA) to validate cell culture conditions. Genetic material was harvested, purified, and purified as described previously. Relative gene expression levels of the gene targets were compared to the housekeeping gene 60 S ribosomal protein L19. Expression levels of all genes were evaluated using the Fast SYBR Green Mastermix (Life Technologies, Grand Island, NY) and custom DNA primers (Integrated, Coralville, IA) in triplicate for three biological replicates.

### Collagen turbidity assay

To evaluate the effect of the PAA/PASP only and HC-PILP on collagen fibrillogenesis, a collagen turbidity assay was performed following previous protocols^35^. Briefly, a 1 mg/ml stock solution of collagen type-I (Thermo Fischer Scientific, Waltham, MA, USA) was prepared by diluting high concentration collagen type-I with PBS and then subsequently kept on ice along with a 96-well plate. Varying concentrations of PAA/PASP only or HC-PILP (Table 1) were then prepared from the stock by adding 0.6-30 μL of the solution to H_2_O. Following dilutions of the HC-PILP and PAA/PASP only stocks, a 1:1 ratio of 1 mg/ml collagen type I and HC-PILP or PAA/PASP only stocks were added to each well resulting in PAA and PASP doses of 0.0234 - 1.18 g and 0.00781 – 0.390 g, respectively. Absorbance measurements at 405 nm were taken every 5 minutes over the span of 2 h at room temperature using a plate reader. The assay also included blank controls of water, PBS, and the highest concentrations of PAA/PASP or HC-PILP.

**Table 1.** Final polymer masses. Volume of HC-PILP nanoclusters or PAA/PASP added into 1 ml of H2O and the final mass of polymer in stock.

| Volume from HC-PILP and PAA/PASP (μL) | Poly Acrylic Acid amount (mg)<br>(MW: 450,000 kDa)<br>Synthesized conc. (39.14 mg/ml) | Poly Aspartic Acid amount (mg)<br>(MW: 14 kDa)<br>Synthesized conc. (13.01) mg/ml |
| --- | --- | --- |
| 30 (High) | 1.18 g | 0.390 g |
| 15 | 0.587 g | 0.195 g |
| 7.5 | 0.293 g | 0.0976 g |
| 3.75 | 0.147 g | 0.0488 g |
| 0.6 (Low) | 0.0234 g | 0.00781 g |

### Attenuated total reflectance Fourier transform infrared spectroscopy (ATR-FTIR)

To determine the protein binding characteristics, ATR-FTIR was performed. Samples were prepared as described in the *Collagen Turbidity Assay*. Samples were incubated at room temperature overnight, immediately flash frozen in liquid nitrogen, and lyophilized. As described previously, samples were background and ATR corrected. Additionally, samples were normalized to the highest ordinate value within the Amide I-II in the fingerprint region in all samples (Amide II was the highest ordinate in all samples)^36,37^. Processed spectral data were transferred to MatLAB for analysis. Spectra were baseline adjusted and the integrated area of the Amide I (1595-1720 cm^-1^) band was calculated. Peak heights were also measured at 1660 cm^-1^ and 1690 cm^-1^. The ratio of 1660 cm^−1^ to 1690 cm^−1^ represents the proportion of non-reducible to reducible cross-links in the collagen, indicative of collagen maturity^38^.

## Statistical Analysis

Data is represented as the mean of biological replicates and error bars represent standard deviation (SD). Measurements were taken from distinct independent samples. Data were analyzed using GraphPad Prism (version 10, GraphPad Software, San Diego, CA). For collagen turbidity and molar relapse data, a repeated measures two-way ANOVA was performed followed by a Tukey’s HSD *post hoc* comparison test to determine statistical difference between independent groups at each respective timepoint. For all other data, a One-way ANOVA was used to determine statistical differences between multiple groups followed by a Tukey’s HSD *post hoc* comparison test. Significant differences were defined at *p* < 0.05. A power analysis was performed prior to the start of animal experimentation with G*Power.

## Results

### Synthesis and characterization of high concentration Polymer-induced liquid precursor nanoclusters (HC-PILP)

For the synthesis of high concentration PILP nanoclusters (HC-PILP), high molecular weight poly acrylic acid (PAA, 450 kDa) and poly aspartic acid (PASP, 14 kDa) were used to chelate calcium (Ca^2+^) at high concentrations. Prior work demonstrated that 9-11 kDA PASP induces rapid and high degree intrafibrillar mineralization^27–29^, we hypothesized that by increasing the MW of PASP it would prevent solution and collagenous structure mineralization. Cryogenic electron micrographs (Cryo-EM) and dynamic light scattering (DLS) exhibit the formation of small ∼2-3 nm structures with no distinct crystalline features (**Fig. 1 A-C**). FTIR spectra of HC-PILP samples exhibit P-O stretching and bending vibrations between 1200-1100 cm^-1^and 700-600 cm^-1^, respectively (**Fig. 1D**). At the P-O bending mode, there is no splitting of the peak indicating a lack of calcium phosphate crystallinity. All these distinct features are absent in the PAA/PASP only samples. To confirm the findings from FTIR, XRD was performed to examine the phase composition (Fig. S1A). In the HC-PILP, some amorphous material was detected prior to pH-adjustment; however, this peak disappeared after increasing the NaOH concentration and only NaCl peaks were detected. The PAA/PASP only samples possess an amorphous diffractogram indicating no crystalline impurities are formed in the synthesis.

**Figure 1.**
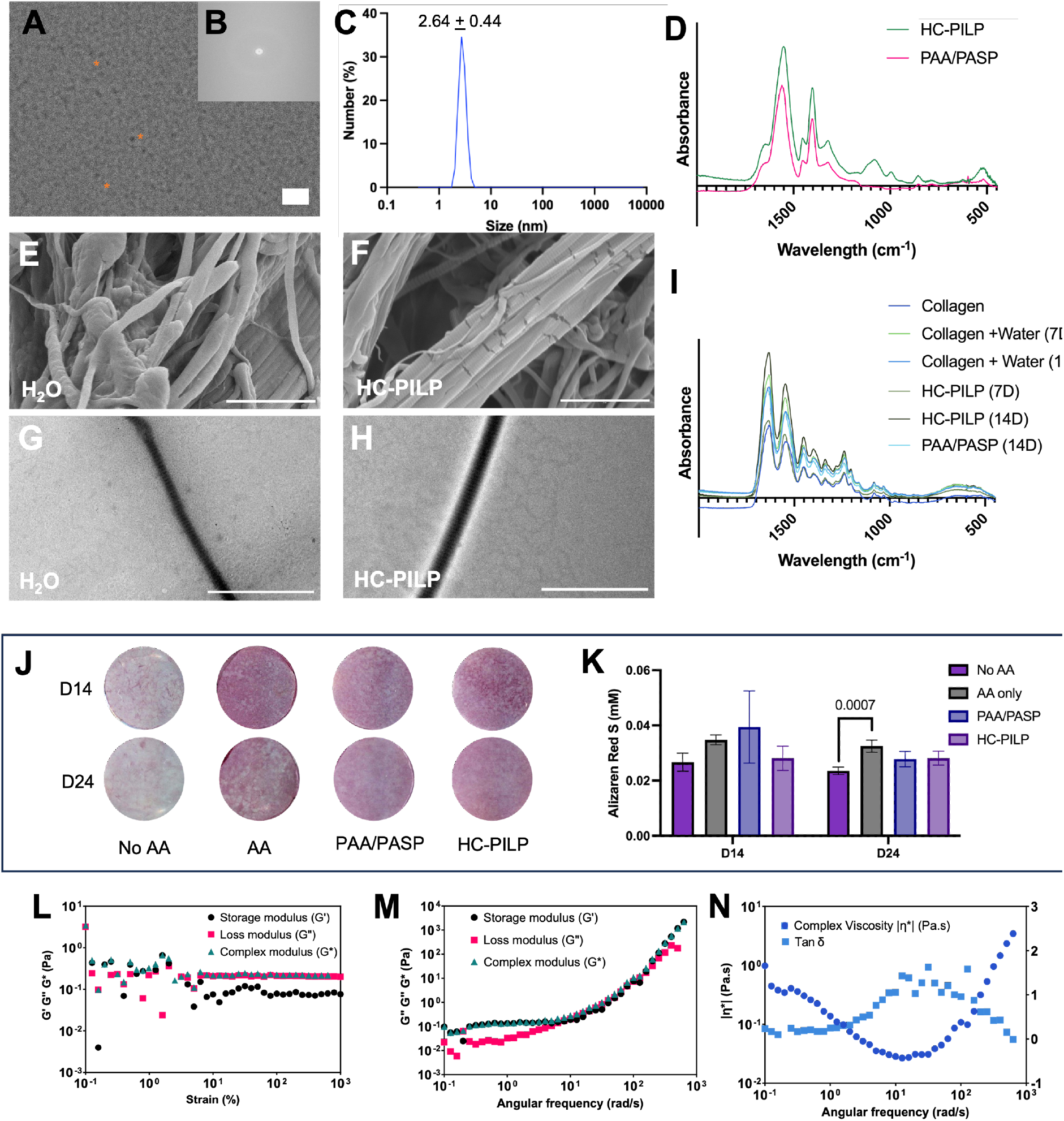
HC-PILP prevents solution precipitation and type-I collagen mineralization. **(A - B)** Cryogenic electron micrograph of densely packed nanoclusters with a corresponding Fourier transform map. Scale bar = 10 nm. **(C)** Dynamic light scattering (DLS) of HC-PILP (n=6 independent synthesis batches). **(D)** Averaged FTIR spectra HC-PILP (green line) and PAA/PASP (pink line) after synthesis (n=3-4 independent synthesis batches). **(E-H**) Scanning and transmission electron micrographs of type-I collagen fibrils after incubation with water (H2O) or HC-PILP for 7 days. Scale bar = 2 μm. **(I)** Averaged FTIR of Type-I collagen fibrils after incubation in H2O, PAA/PASP, HC-PILP for 7 or 14 days (7D or 14D) (n=3-4 independent samples). **(J)** Alizarin red S (red/pinkish dye) of MC3T3-E1 cells. Ascorbic Acid (AA) was used to induced matrix deposition in all groups. Samples were then treated with PAA/PASP or HC-PILP for an additional 14 or 24 days. No AA was used as a negative control. **(K)** Bar graph (mean ± sd) of extracted alizarin red S dye from MC3T3-E1 cells after PAA/PASP and HC-PILP treatment. P<0.05 determined by One-way ANOVA (n=4). (**L-N**) Rheological analysis of HC-PILP.

Next, *in vitro* mineralization experiments were performed on insoluble collagen Type-I. After 7 days of incubation with HC-PILP, extrafibrillar and intrafibrillar mineralization were not observed and the periodicity of the collagen fibrils can be seen (**Fig. 1E-H**). Furthermore, collagen fibrils do not show any distinct phosphate spectral peaks after 7 and 14 days. As controls, collagen was incubated in water and PAA/PASP for 14 days and the spectral patterns are similar to the HC-PILP treated fibrils (**Fig. 1I**). Further validating the lack of extra and intrafibrillar mineralization of insoluble collagen, EDX elemental analysis showed a lack of calcium and phosphate within the HC-PILP treated collagen (Fig. S1B-E). Importantly, the collagen maturity is not altered after incubation in these conditions (Fig. S1F). To examine mineralization in more dynamic settings, an *in vitro* mineralization assay was performed on MC3T3-E1 cells with the calcium-binding dye Alizarin red. Without Ascorbic Acid (AA), the cells exhibit a lack of distinct alizarin-stained calcium nodules at 14 and 24 days (**Fig. 1J**). After 14 and 24 days of incubation with AA, mineralization becomes more apparent and evenly dispersed within the wells containing AA-treated cells. Similar findings were observed for the PAA/PASP and HC-PILP treated samples. Subsequent quantification of the alizarin-bounded dye does not show a significant difference between any conditions at 14 days, but a significant increase after 24 days between the AA-treated and non-AA-treated cells was observed, confirming the qualitative results (**Fig. 1K**). Lastly, the HC-PILP was able to induce an osteogenic phenotype *in vitro* (Fig. S1G-I). Taken together, these results show that the incorporation of 14 kDA PASP leads to the formation of solution stable calcium phosphate nanoclusters that can prevent solution precipitation of calcium phosphate as well as prevent the extrafibrillar and intrafibrillar mineralization of collagen Type-I and cells.

The examination of HC-PILP mechanical properties involved employing strain-dependent and frequency-dependent oscillatory rheology. Across a strain amplitude from 10 to 10k % and a consistent frequency of ∼10 rad/sec (**Fig. 1L**), the storage modulus (G’), loss modulus (G”), and complex modulus (G*) of HC-PILP maintained constancy and independence. This observation delineated an expansive linear viscoelastic region (LVR), indicating the nanoclusters’ resistance against deformations or structural alterations. Moreover, the concurrent instances of G” > G’ and G* aligning with G” displayed HC-PILP as a viscoelastic liquid. Expanding on the LVR, two distinct crossover frequencies were observed at ∼10 rad/sec and ∼100 rad/sec with constant strain and varied oscillatory frequencies (**Fig. 1M**). Below and above the crossover frequencies, the material exhibited predominantly viscoelastic solid behavior (G’ > G’)’, signifying a more elastic response capable of recovering upon deformation. Conversely, within the range of the two flow-like demeanor devoid of significant elastic recovery. The complex viscosity |η*| and Tan δ curves exhibited two crossover frequencies confirming the complex rheological nature of HC-PILP characterized by multiple relaxation processes (**Fig. 1N**). These transitions between elastic and viscous behavior at the crossover frequencies delineated the intricate interplay of diverse relaxation mechanisms, exemplifying the material’s capability to shift between solid-like and fluid-like responses under varying deformation frequencies. Hence, HC-PILP exhibited characteristics of shear-jammed colloidal material, confirming the formation of nanoclusters.

### HC-PILP minimizes orthodontic relapse in a rat model

Since results show that a higher PASP molecular weight is able to prevent the solution precipitation of calcium phosphate, minimize extrafibrillar and intrafibrillar mineralization, and induce an osteogenic phenotype in vitro, studies were conducted to determine if HC-PILP would minimize orthodontic relapse by improving the underlying bone without inducing ectopic mineralization of the PDL. To test this hypothesis, Sprague Dawley rats were used to induce mesial movement of the left maxillary 1^st^ molar (**Fig. 2A-B**). The final amount of tooth movement after 28 days showed no statistical difference (**Fig. 2C**). In addition, the weight of the animals did not change throughout the duration of the experimental timeline indicating the experimental groups did not alter the feeding habits on the animals (Fig. S2A). Moreover, HC-PILP exhibited a significant reduction of relapse over 24 days relative to the associated controls (**Fig. 2D** and Fig. S2B-F). After 4 days of relapse, HC-PILP and PAA/PASP significantly lowered the percentage relapse of the initial tooth movement (21.74 + 14.89% and 25.14 + 8.27%, respectively) contrasting with the PBS relapse (40.0 + 12.97%). HC-PILP further significantly reduced the amount of relapse after 12 and 24 days of relapse (32.58 + 14.88% and 55.99 + 13.49%, respectively) compared to both PAA/PASP (52.11 + 8.27% and 81.33 + 22.47%) and PBS (52.75 + 4.87% and 78.99 + 14.13%). Next, *ex vivo* μCT was performed to determine if the underlying interradicular bone improved during the relapse phase. No differences were observed in the bone volume fraction (BVF) and bone mineral density measurements (BMD) after 24 days of relapse between the experimental groups (Supplemental Table 1). The microarchitecture of the trabecular bone shows a gradual improvement after HC-PILP administration after 12 days of relapse. Lastly, the PDL space is non-mineralized as assessed from the μCT scans (Fig. S2B-F). These findings indicate that the bone density does not appear to improve, but the trabecular microarchitecture does after HC-PILP treatment later in relapse and the treatment does not induce ectopic mineralization of the PDL space.

**Figure 2.**
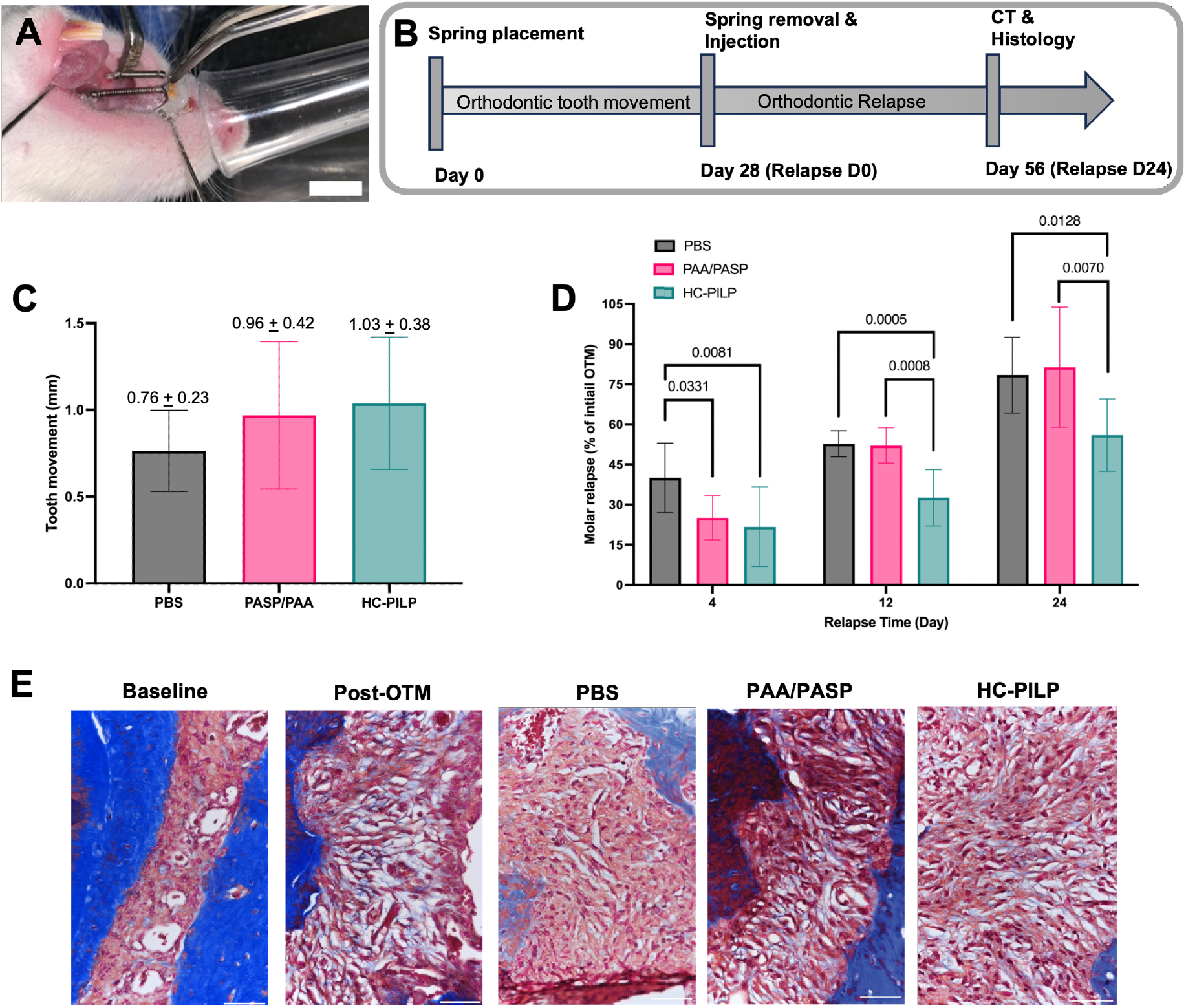
HC-PILP treatment minimizes relapse in a rat model of orthodontic relapse. **(A)** 12-week-old Sprague Dawley rat with a 12 mm closed-coil spring attached to the maxillary mesial-buccal surface of the 1^st^ molar and the ipsilateral surface of the incisor. Scale bar = 10 mm. **(B)** Experimental timeline of inducing orthodontic tooth movement for 28 days, which is followed by removal of the springs and a single injection of PBS, PAA/PASP, or HC-PILP in 30 μL. Relapse is assessed for an additional 24 days with subsequent micro-CT analysis and histology. **(C)** Bar graph (mean + sd) of the first 28 days of tooth movement from PBS (Black), PAA/PASP (Pink), and HC-PILP (Green). P<0.05 determined by One-way ANOVA (n=14-17). **(D)** Molar distal relapse presented as a percentage of initial molar movement after a single injection of PBS (Black), PAA/PASP (Pink), and HC-PILP (Green). p<0.05 determined by Two-way ANOVA with a Tukey’s post-hoc test for multiple comparisons (n=4-7, per group at each time point). All data is represented as mean and standard deviation. **(E**) Masson Trichrome micrographs (40x) at the mesial surface of the distal buccal root of baseline and post-orthodontic tooth movement (Post-OTM) controls as well as PBS, PAA/PASP, and HC-PILP at 4 days relapse. Scale bar = 50 μM.

### HC-PILP alters PDL remodeling during early relapse

Although a modest improvement in the interradicular trabecular bone was seen later in the relapse, no significant bone alterations were seen in the early stages. Based on these findings, the PDL was further investigated given its implication in relapse. Here, the mesial surface of the distal buccal root, which represents the compression side during OTM, was examined (**Fig. 2E** and Fig.S3). Prior to tooth movement, the collagen fibers fully span the entire PDL space with high cellularity and blood vessel presence. Upon removal of the orthodontic appliances, the post-OTM PDL compression-side appears less organized with discontinuous fiber orientation and a distribution of red blood cells throughout the space. After 4 days of relapse, the abundance of collagen, cellular infiltration, and organization increases in the PBS group. PAA/PASP treatment exhibited slightly less recovery than the PBS group with some reorganization of the PDL space. HC-PILP exhibited a similar phenotype to the post-OTM with sparse collagen abundance, disorganization of the fibers, and a lack of cellular components. Over the remainder of the relapse phase, the PDL cellular infiltration increases at relapse day 12 in all the groups; however, the collagen distribution of the HC-PILP treated rats still was dampened. By day 24 of relapse, the PDL integrity returned to baseline levels (Fig. S4 and S5).

These findings suggest a disruption of PDL remodeling after intervention with HC-PILP. To further investigate the collagen content and maturity within the PDL, picrosirius red staining with polarized light microscopy was performed. Picrosirius red is a collagen-sensitive dye that allows for distinguishing collagen orientation and its subsequent maturity (**Fig. 3A**)^39^. Like the Masson trichrome stain, collagen content was significantly reduced and was mostly absent post-OTM (**Fig. 3B**). After PBS administration, collagen content showed an increase at relapse day 4 and full recovery to baseline by day 12 (**Fig. 3C-D**). The orientation of the fibers spans the entire PDL, and the number of fibers covers the entire PDL space by relapse day 24 (**Fig. 3E**). After PAA/PASP treatment, the collagen content showed some similar recovery with fibers spanning the entire PDL by relapse day 24 (**Fig. 3F-H**). However, the orientation appears to be altered until relapse day 24. Strikingly, after HC-PILP administration, collagen content is significantly reduced after 4 and 12 days of relapse (**Fig. 3I-J**). There is minimal signal observed at early relapse day 4. After 12 days, some collagen signals appear but do not span the entire PDL, unlike PBS and PAA/PASP, which fully recovered to similar content levels as the baseline. After 24 days, the collagen abundance improved, but the orientation slightly differed from the baseline control **(Fig. 3K**). To validate these results, the signals of independent color fibers were quantified. The red fibers of the HC-PILP group were at lower levels similar to the post-OTM and significantly lower than the PBS and PAA/PASP groups at 4 days of relapse (**Fig. 3L**). After 12 days of relapse, the PAA/PASP and PBS groups returned to similar levels as the baseline, but HC-PILP remained significantly lower than the PBS group. Lastly, after 24 days of relapse, all groups’ red fiber content was higher than the baseline but was not statistically different. Similar trends were observed for the green fibers in that PILP treatment reduced collagen detection during relapse (**Fig. 3M**). Additionally, the distal surface of the mesial root, which represents the tension side during OTM, was examined. In contrast to the observations for the compression side, the mesial root undergoing tension displayed less collagenous remodeling after PILP treatment (Fig. S5-7). Importantly, the collagen within the PDL space recovered with no differences observed from the baseline controls. Overall, these findings show that HC-PILPs are able to modify collagen remodeling within the PDL space, particularly on the compression side surface after relapse, and the amount of relapse post-OTM at early stages.

**Figure 3.**
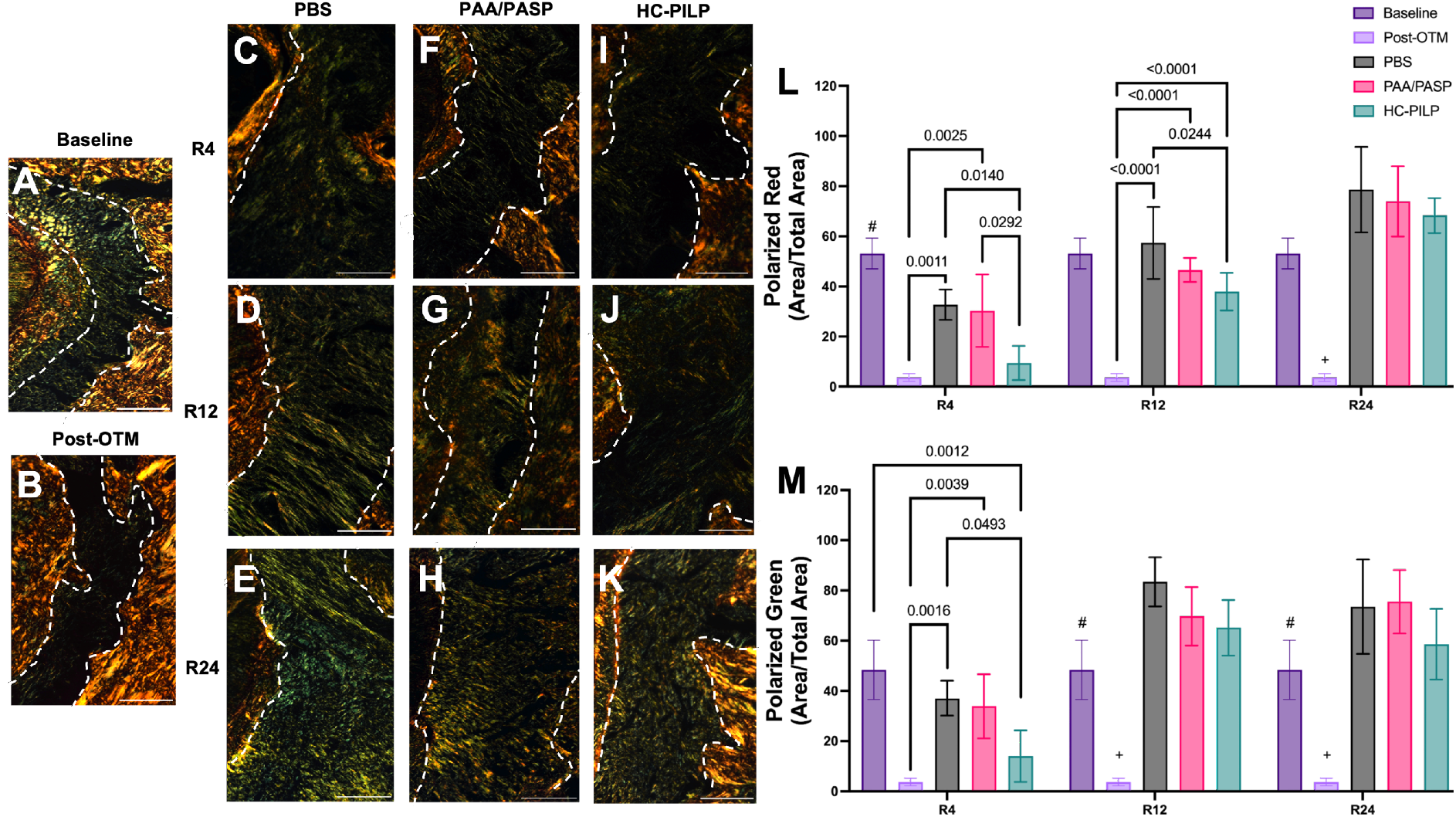
HC-PILP treatment alters periodontal ligament remodeling during relapse. **(A-K)** Picrosirius polarized light microscopy at the mesial surface of the distal buccal root. Representative images are shown after treatment with PBS **(C-E)**, PAA/PASP **(F-H)**, and HC-PILP **(I-K)** after 4, 12, and 24 days of relapse. Baseline and Post-OTM are shown for comparison. White dashes outline indicates bone and tooth borders. Scale bar = 50 μM. Bar graph of collagen detection across different color channels: Red **(L)** and Green **(M)**. p<0.05 determined by One-way ANOVA with a Tukey’s post-hoc test for multiple comparisons (n=3-6, per group at each time point). All data is represented as mean and standard deviation.

### HC-PILP modulate collagen fibrillogenesis and protein secondary structure

Given that PDL remodeling was altered after HC-PILP administration, the potential mechanism by which HC-PILP induced this response was investigated. First, the question of whether this response was influenced by cellular behavior or cells’ ability to make collagen was addressed. To test this hypothesis, NIH-3T3 fibroblasts were stimulated for 48 hours (**Fig. 4A**). After HC-PILP administration, gene expression of *Col1* and *MMP-2* was not altered. A 2-fold upregulation was observed for the PAA/PASP only control, but this was not significantly different from the non-treated control. Upregulation of alpha-smooth muscle actin (α-SMA) was observed after PILP and PAA/PASP treatment similar to the treatment with TGF-β (Fig. S8). From the gene expression data, no distinct changes were observed after HC-PILP treatment suggesting an alternative mechanism from the initial hypothesis.

**Figure 4.**
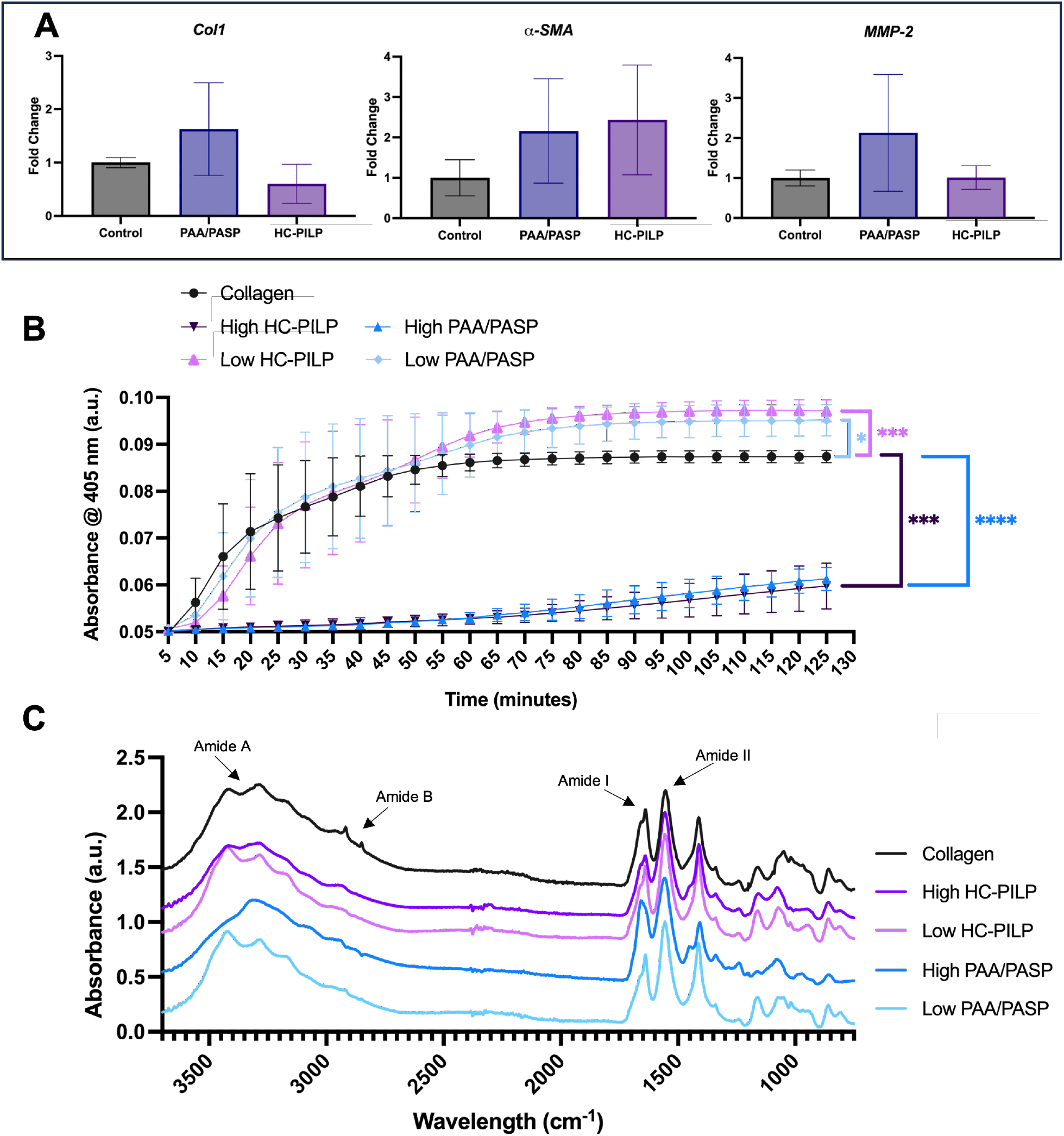
HC-PILP treatment and PAA/PASP alter fibrillogenesis and protein secondary structure. (**A**) Bar graph of gene expression data from NIH-3T3 fibroblast. (**B)** Type-I collagen fibril formation detected with absorbance (405 nm) over 120 minutes. **** indicates p<0.0001, *** p<0.0005, and * p<0.05 determined by One-way ANOVA.(**C**). FTIR Spectra of Type-1 collagen after firbillogenesis in the presence of High HC-PILP (purple), Low HC-PILP (pink), High PAA/PASP (dark blue), and low PAA/PASP (light blue).

Collagen molecules must assemble to form fibrils and there is a possibility that this process may be impacted by HC-PILP. A high and low dose of HC-PILP and PAA/PASP were chosen to be in the presence of collagen molecules during fibrillogenesis (**Table 1** and **Fig. 4B**). Fibril formation kinetics were significantly impacted and dependent on the HC-PILP concentration. The high dose HC-PILP yielded the greatest impact at 120 min. Interestingly, the PAA/PASP also had a clear effect on fibril formation and diminished fibrillogenesis. As the dose was reduced, a clear dose-response was observed for both groups (Fig. S9). However, at the low concentration, fibril formation was enhanced in the presence of both PAA/PASP and HC-PILP. Fibril formation kinetics were slightly slowed and then elevated higher than the collagen control. Based on these findings, we wanted to assess if any changes could be observed in the secondary structure of the collagen type-I fibrils. FTIR was performed after a 24-ahour incubation of collagen molecules with high and low doses of both HC-PILP and PAA/PASP. Collagen alone exhibits distinct spectral bands corresponding to Amide A at 3400-3100 cm^-1^ (O-H and N-H stretching) and Amide B at 3000-2900 cm^-1^. In the fingerprint region, Amide I was observed at 1639 cm^-1^ (corresponding to carbonyl (C=O) stretching). Amide II at 1554 cm^-1^ and is more complex than Amide I (mainly due N-H bond bending and C-N and C-C bending), and Amide III at 1300-1200 cm (C-N stretching and N-H deformation). Additional peaks were identified between 1460 cm^-1^ and 1417 cm^-1^ corresponding to CH2 bending. In the presence of HC-PILP, a bathochromic shift (left shift) of the Amide I peak occurred with a significant reduction in the area under the curve (AUC) peak (**Fig. 5** and **Table 2**). A similar bathochromic shift was observed for the PAA/PASP treated collagen fibrils; however, no difference was observed for the AUC of Amide I. The low-dose PAA/PASP and HC-PILP exhibit the same spectral patterns with Amide I peak heights. Amide II and Amide III did not exhibit any bathochromic or hypsochromic shifts in any of the groups. These results show that high HC-PILP and PAA/PASP can modulate collagen fibrillogenesis and protein secondary structure albeit in different manners, but low doses do not affect protein structure or elevate collagen type I formation.

**Table 2.** Summary of Amide I area of under the curve (AUC) and peak wavenumber between 1700-1600 cm^-1^. a= p< 0.05 vs. collagen and b= p< 0.5 vs High PAA/PASP.

| Condition | AUC Amide I | Peak wavenumber |
| --- | --- | --- |
| Collagen | 22.35 ± 1.70<sup>a</sup> | 1639 |
| High HC-PILP | 11.49 ± 4.19<sup>a, b</sup> | 1647 |
| Low HC-PILP | 17.72 ± 3.17 | 1639 |
| High PAA/PASP | 20.47 ± 2.78<sup>b</sup> | 1658 |
| Low PAA/PASP | 15.27 ± 0.81 | 1639 |

**Figure 5.**
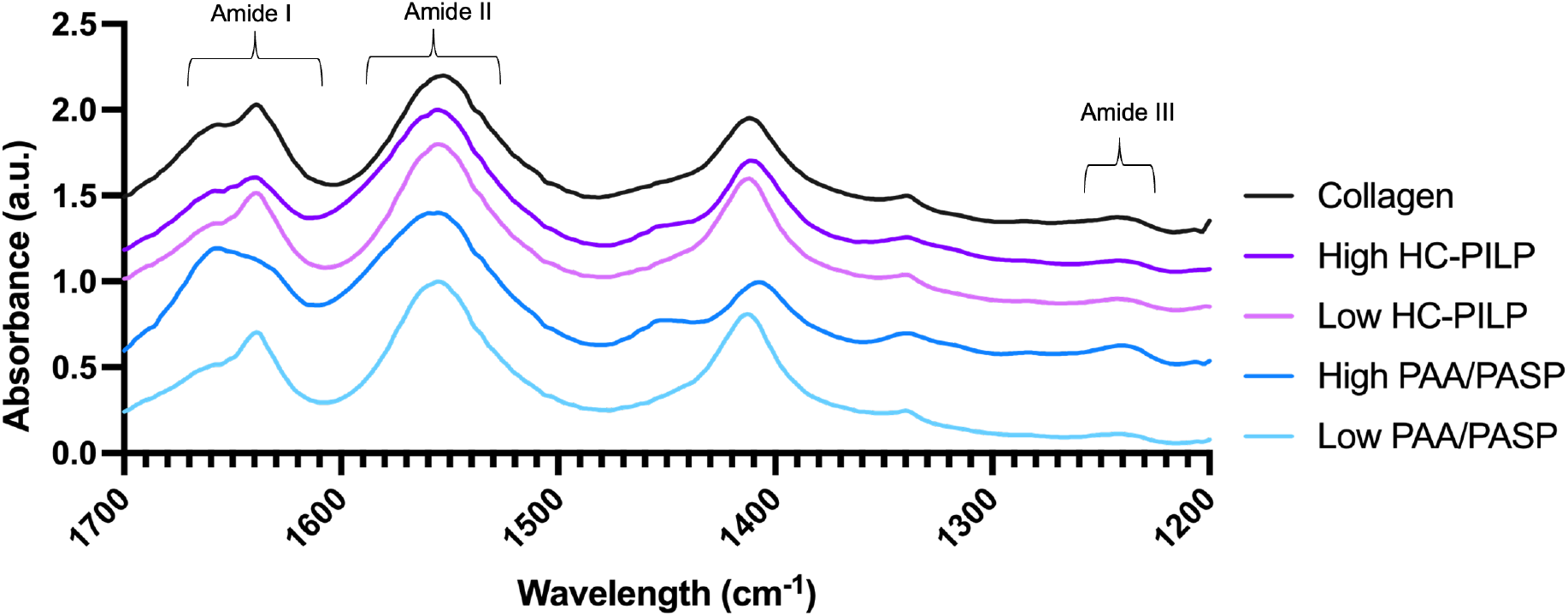
HC-PILP treatment and PAA/PASP alter fibrillogenesis and protein secondary structure. FTIR Amide region of Type-1 collagen after firbillogenesis in the presence of High HC-PILP (purple), Low HC-PILP (pink), High PAA/PASP (dark blue), and low PAA/PASP (light blue).

## Discussion

Orthodontic relapse is a critical issue occurring after orthodontic therapy and current modes of tooth retention are still unable to provide adequate stability. Current retention methods include non-surgical approaches with removable- and fixed-retainers and surgical interventions by cutting the attaching PDL to minimize relapse^8^. Nonetheless, both methods still have a high probability of orthodontic relapse and new methods to minimize relapse outcomes are desired. Here, an approach to minimize orthodontic relapse by local delivery of HC-PILP is discussed.

The periodontium is a complex interphase involving the non-mineralized PDL attaching the tooth to the underlying bone. When aiming to target the periodontium for any therapy it is important to minimize the risk of mineralizing the PDL, a process known as ankylosis, while also improving or altering the tooth supporting structures^40^. To achieve this, a HC-PILP method was utilized. HC-PILPs have been shown to facilitate mineralization and biological responses to induce bone formation by combining PASP and PAA together. Interestingly, the concentrations of HC-PILP polymer, calcium, and phosphate are much higher than in traditional PILP methods^27–29^.In these modified HC-PILP formulations, potent mineralization of collagen type-I is achieved when using 9-11 kDa PASP. Here, a high chain length PASP was used given that other studies were able to stabilize solution calcium and phosphate as well as prevent solution precipitation, albeit lower concentrations^30^. 14 kDa PASP was incorporated into the HC-PILP solution, and the nanoclusters were stable in solution. CryoEM exhibits dense liquid nanoclusters with even homogeneity and dispersion in solution. However, after liquid removal via lyophilization, no amorphous structures were observed after the pH-adjustment. This is contrary to works from Yao et al. that describe similar dense structures with amorphous phases in the remaining dried material. These differing findings could primarily be attributed to the increased chain length of PASP. High MW PASP on its own can impede solution precipitation^30,41^. The commercially available PASP utilized in this study has been demonstrated to have a disordered secondary structure and this may facilitate increased flexibility and its ability to chelate additional ions in solution to prevent solution precipitation than lower MW PASP^41^. With the addition of PAA, another strong calcium chelator^42^, solution precipitation and fibril mineralization are fully inhibited. This could be because the combination of polymers stabilizes the nanoclusters against colloidal aggregation and preventing mineralization^43^. These results were further validated when used in the presence of insoluble collagen type-I and osteoblastic cell lines for *in vitro* mineralization. This supports works from others suggesting concentrations higher than 1 mg/ml of polyanionic prevent mineralization^24^. Interestingly, osteogenic effects were maintained *in* vitro despite the lack of mineralization induction. Taken together, the incorporation of high MW PASP into modified HC-PILPs minimizes solution precipitation and inhibits collagen mineralization. These features are ideal when aiming to maintain high concentrations of calcium and phosphate in solution for local periodontal therapy. Importantly, HC-PILPs synthesized in this study are different from traditional PILP methodology and do not induce typical mineralization but maintain similar *in vitro* osteogenic effects. These results add additional insight into modifications that can alter the mineralization and biological behavior of HC-PILPs.

Orthodontic relapse biology is complex and thought to proceed through reversal of the original biological reactions that occurred during the initial OTM. This includes bone loss adjacent to the relapsing tooth and remodeling of the PDL^12,14^. To mitigate this, various groups have focused on the use of drugs and other biological approaches to minimize relapse. An alternative approach would be the use of biomaterials like calcium and phosphate given their precise synthesis methods and ability to be injected^40^. Importantly, previous studies have not been able to minimize relapse alone without any drugs or morphogenic additives. We hypothesized that HC-PILP would improve the bone quality of the underlying bone and subsequently improve relapse. However, this hypothesis was only true for late-stage relapse. The μCT data shows that the bone density did not improve relative to the post-OTM controls. This correlates well with other works on osteoporotic bone and HC-PILP treatment, where bone density improvements were not observed as early as 4 weeks^27,29^. Importantly, HC-PILP showed improvements in the trabecular bone. Although a modest improvement in bone was observed, it did not provide evidence for improving early-stage relapse.

Collagen remodeling within the PDL has been strongly associated with early-stage relapse. Results show that HC-PILP significantly alters the collagen remodeling in the PDL after 4 days of relapse. This effect was maintained for as long as 12 days prior to the PDL integrity returning to baseline levels after 24 days of relapse. The dampened PDL remodeling correlated with a decrease in the percentage of relapse. Previous works demonstrated that pressure forces degraded the PDL collagen fibers, and its rapid recovery corresponded with an increase relapse. Altering PDL recovery with TGF-β and relaxin supports the hypothesis that early-stage relapse is associated with collagen recovery within the PDL^12,13^. The findings in this study support other groups’ work showing that relapse can be mitigated by modulating PDL remodeling.

Collagen formation is a self-assembling process onset by the aggregation of collagen monomer as nuclei that will grow and assemble into fibrils^44^. A major finding of this work is the effect of collagen type-I fibrillogenesis after exposure to the polymers used in HC-PILP synthesis. Studies describing PILPs have focused on collagen mineralization and how mineralization is affected by the collagen cross-linking mechanism^45,46^. No study has been reported on the occurrence of collagen type-I fibrillogenesis in the presence of PILP and the associated polymers for PILP synthesis. Non-collagenous proteins (NCPs) are soluble acidic proteins that appear in the composition of bone and play key roles in signaling and matrix mineralization^47^. PASP and PAA utilized in the HC-PILP synthesis have been demonstrated to emulate NCP mineralization^25,26^. More recently, Depalle et al. described the role of Osteopontin, a well-known NCP, and its role in collagen type-I assembly and matrix structure^48^. Results described here provide additional evidence that acidic polymers (i.e., PASP and PAA) affect collagen fibril formation. Collagen molecules were incubated directly with the polymers and this altered the fibril formation as assessed with absorbance, suggesting an intimate interaction between collagen molecules and the polymers. To examine the potential mechanism by which this occurred, additional FTIR was performed to examine the protein secondary structure. Amide I is considered a sensitive marker for alterations in the helical polypeptide backbone^49^. Results show that there are distinct bathochromic shifts from normal collagen (1639 cm^-1^) to 1647 and 1658 after fibril formation in the presence of HC-PILP and PAA/PASP, respectively. This may indicate a change in the collagen conformation. This may be due to potential hydrogen bonding interactions and/or electrostatics influencing collagen assembly, which is supported by the higher wavenumber shift. The shift suggests weakened molecular bonds between the collagen molecules and potential bonds forming between collagen and the polymers. Interestingly, the area under the curve (AUC) of Amide I was significantly different from the collagen control and HC-PILP treated collagen, which was not observed for other conditions. This difference can be attributed to HC-PILP formed in our synthesis. Liquid-liquid phase separation occurs when calcium and phosphate are in solution, and this may provide more stable structures to interact with specific regions on collagen molecules and impede collagen formation at high doses^50^.

Surprisingly, low doses of HC-PILP and PAA/PASP enhanced fibril formation relative to the collagen control. No spectral differences were observed between the low dose treated groups and collagen alone. Within the first 20 minutes of fibrillogenesis, fibril kinetics were slower than the collagen control. However, after an hour fibrillogenesis was elevated. The PILP polymers may be promoting growth of the fibrils. As seen with the high dose, the PILP polymers can interact and impede collagen assembly by effecting bond formation. Lower doses may promote similar interactions; however, these bound structures create sites for collagen aggregation and enhances the growth phase of collagen fibrils. Similar findings have been observed for other materials and their ability to promote collagen nuclei formation and subsequent fibril formation^49^.

While our results are promising, we recognize that there may exist some limitations to the work described. The animal studies utilize high orthodontic spring forces and this causes significant bone loss. Although it is ideal to get adequate tooth movement within 28-day OTM, less forces may be more clinically relevant. In addition, only male Sprague-Dawley rats were utilized in this study. The use of male rats is a common theme in the orthodontic biology field, but the assessment of female rats is necessary for future studies to examine therapeutic outcomes and to assess sex as a biological determinant. Lastly, mucosal injections are technique-sensitive, and the total volume of injection solutions may be altered. This may have been minimized by the viscoelastic behavior of the HC-PILP material. The elastic component (G’) at both high and low frequencies, above and below the two crossover frequencies, respectively, may facilitate material conformity to the tissue surface, resistance against deformation due to relapse recoil, masticatory forces, and maintenance of material integrity at the injection site. Moreover, the dominant viscous component (G’’) within the two transition points promises enhancements in material spreading, integration, and tissue residence time post-injection. Thus, the rheological characterization properties showed the suitability of supportive mechanical attributes and validated HC-PILP as a suitable injectable biomaterial for mucosal injections^51^. Additionally, while power analysis shows that we were able to achieve significant power in the current study, future work will benefit from larger sample sizes.

In this work, a method to treat orthodontic relapse by altering the periodontal tissues at early and late stages of relapse using a biomaterial approach is described. The use of HC-PILP and their ability to alter PDL remodeling during early relapse and improve bone quality at later stages was demonstrated. In addition, the use of different PASP molecular weights to control mineralization, as well as the interaction between acidic polymeric analogs and their ability to alter collagen fibril formation was explored. This work provides a promising approach to treating orthodontic relapse without the need for other morphogenic factors. Applications of this nanocluster platform in additional clinically relevant models, including joint enthesis, fibrotic tissue, and endochondral fracture represent existing prospects to further advance our ability to modulate tissue repair.

## Contributions

D.L.C., S.H., S.D.K., and T.A.D. conceptualized the project and designed the experiments. D.L.C. performed the material characterization and animal experiments. D.L.C., M.L.F., and J.C., contributed to stone model fabrication, histology, and cellular data generation. B.N.K. characterized rheological properties of the material. G.J.K provided the MatLab script for FTIR analysis. A.J. provided critical feedback on experiments throughout the study. D.L.C., S.H., S.D.K., and T.A.D. interpreted the results. D.L.C. wrote the first draft of the study and all subsequent versions were edited and approved by the authors. D.L.C., S.D.K., and T.A.D. received fundings. All authors listed have made a substantial, direct, and intellectual contribution to the work and approved it for publication.

## Acknowledgements

We would like to thank Nick Szeto and Wenhan Chang at UCSF’s Core Center for Musculoskeletal Biology and Medicine for microCT analysis, David Buckley at the UCSF Bay Area cryo-EM consortium for assistance with Cryo-electron micrographs (Cryo-EM equipment at UCSF is partially supported by NIH grant S10OD020054 and Howard Huges Medical Institute), and Nick Settineri for X-ray diffraction assistance in the Small Molecule X-ray Crystallaography Facility at UC Berkeley (NIH Shared Instrumentation Grant S10-RR027172) Lastly, we would like to thank Preethi Rhagavan for her thoughtful feedback on the manuscript.

