## Supplementary figures and images for "Calcium phosphate nanoclusters modify periodontium remodeling and minimize orthodontic relapse"

### Supplemental Figures 1-9

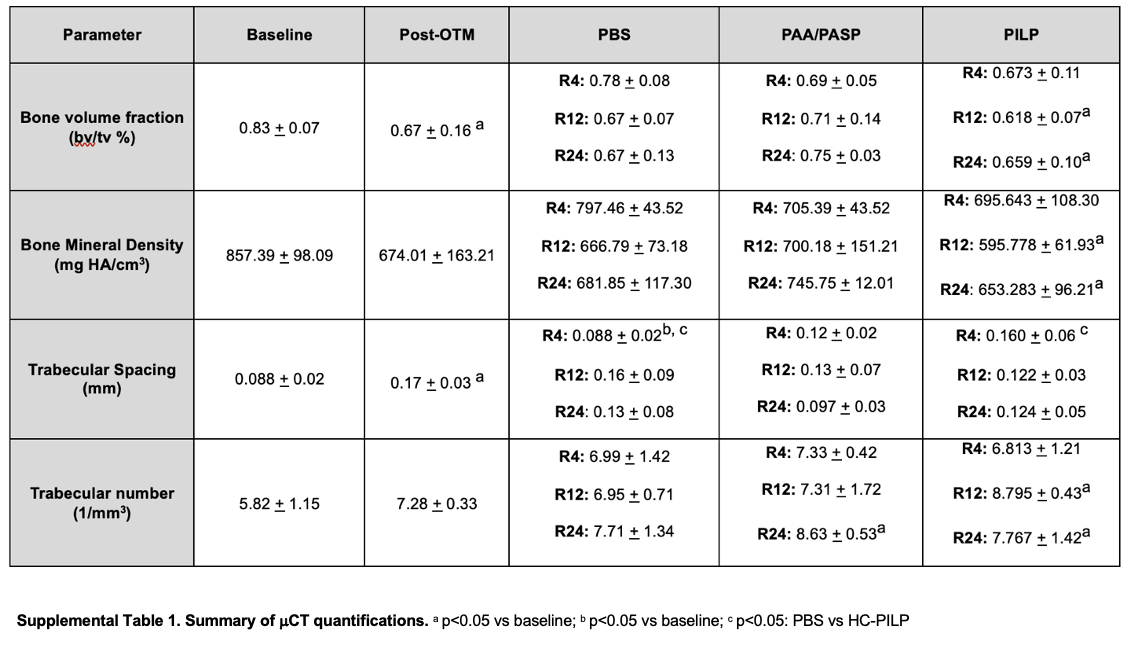

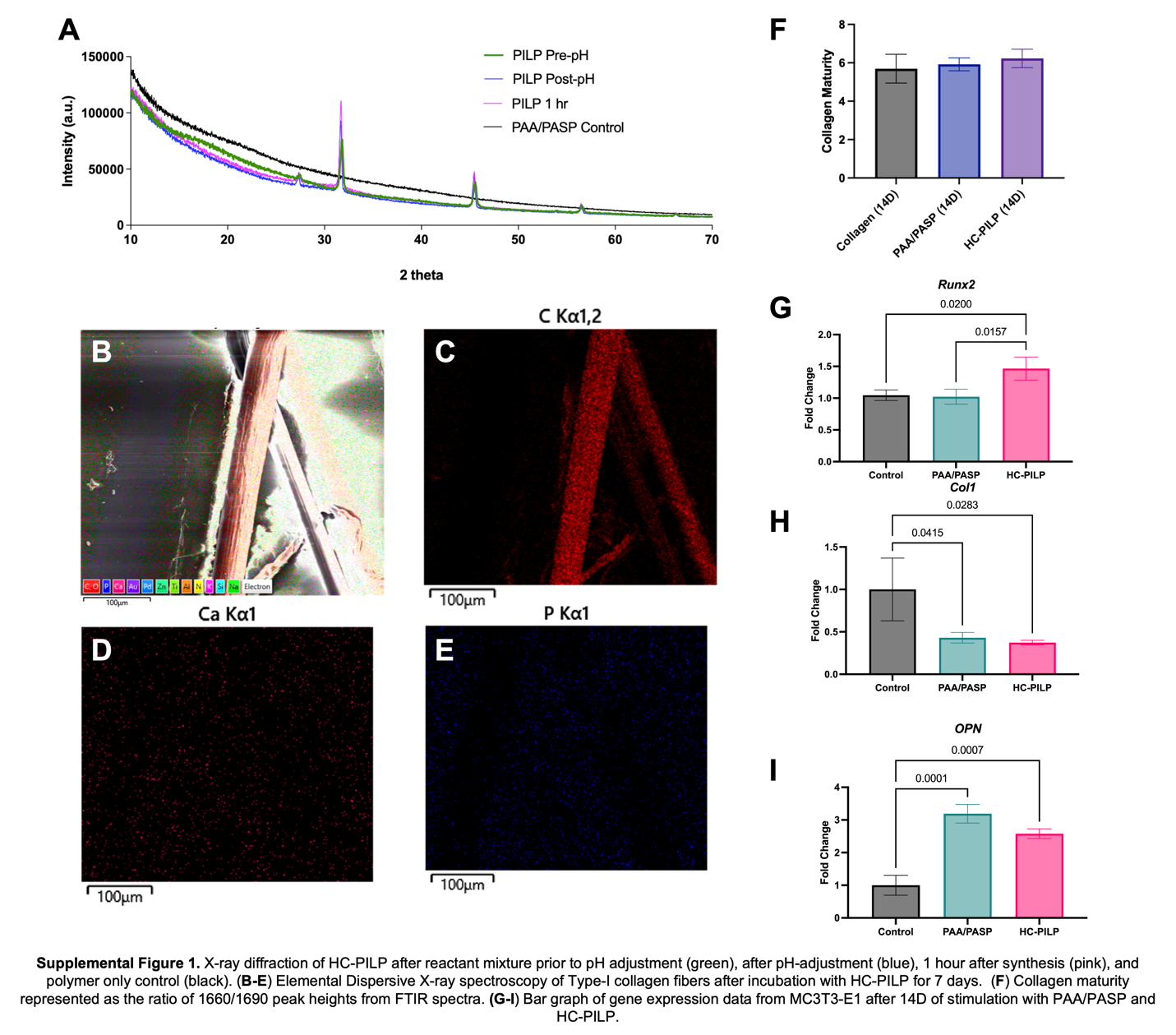

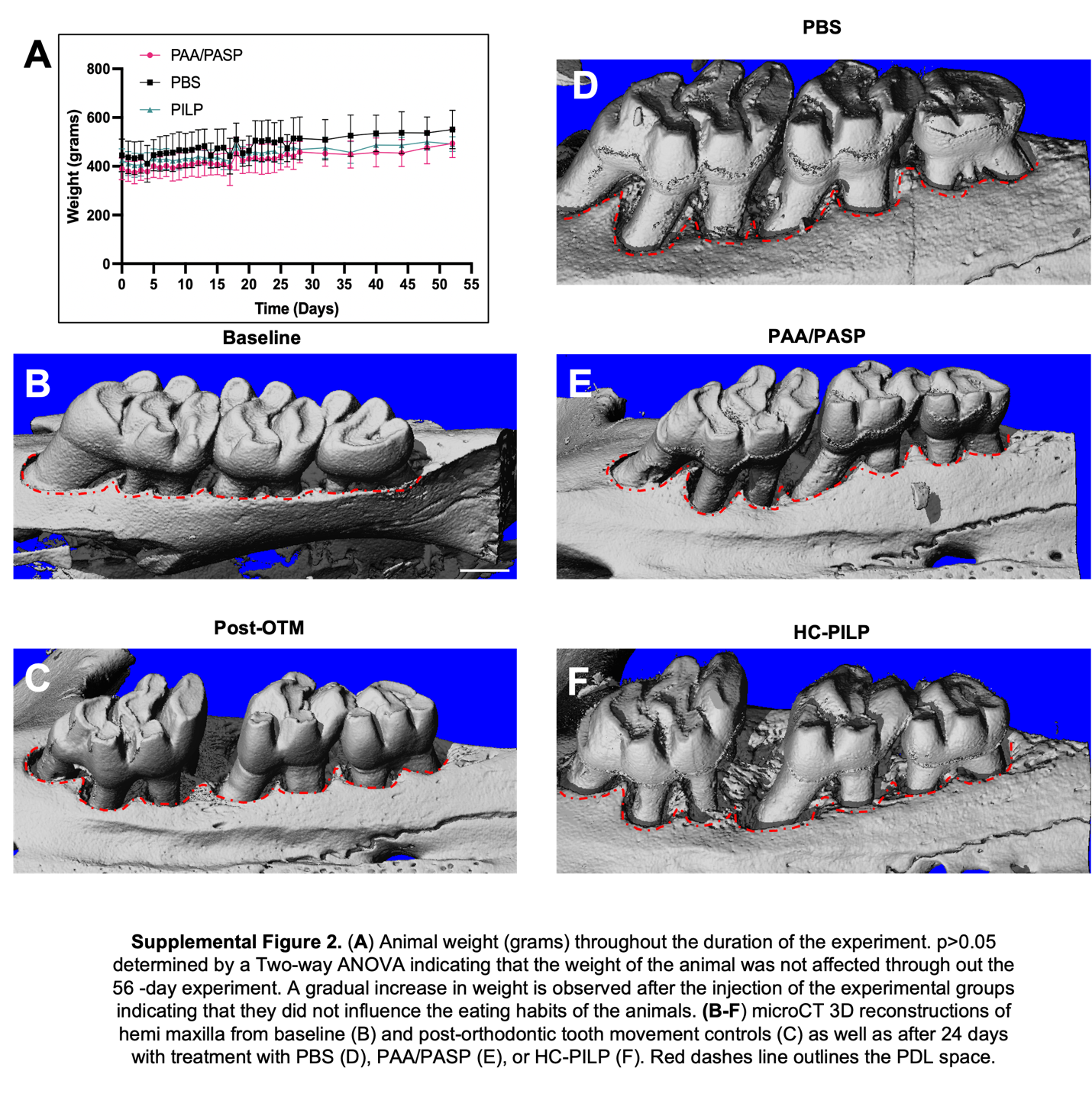

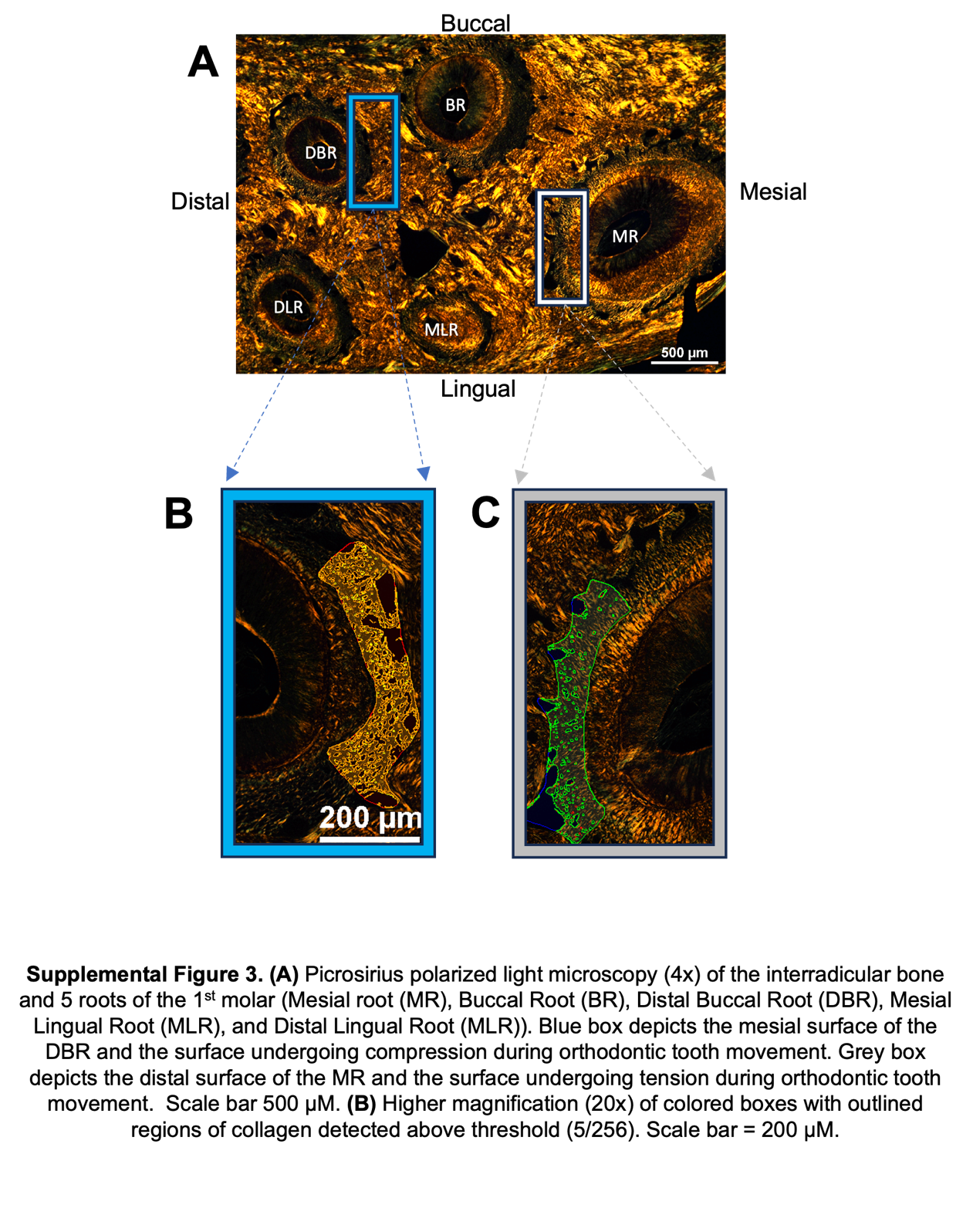

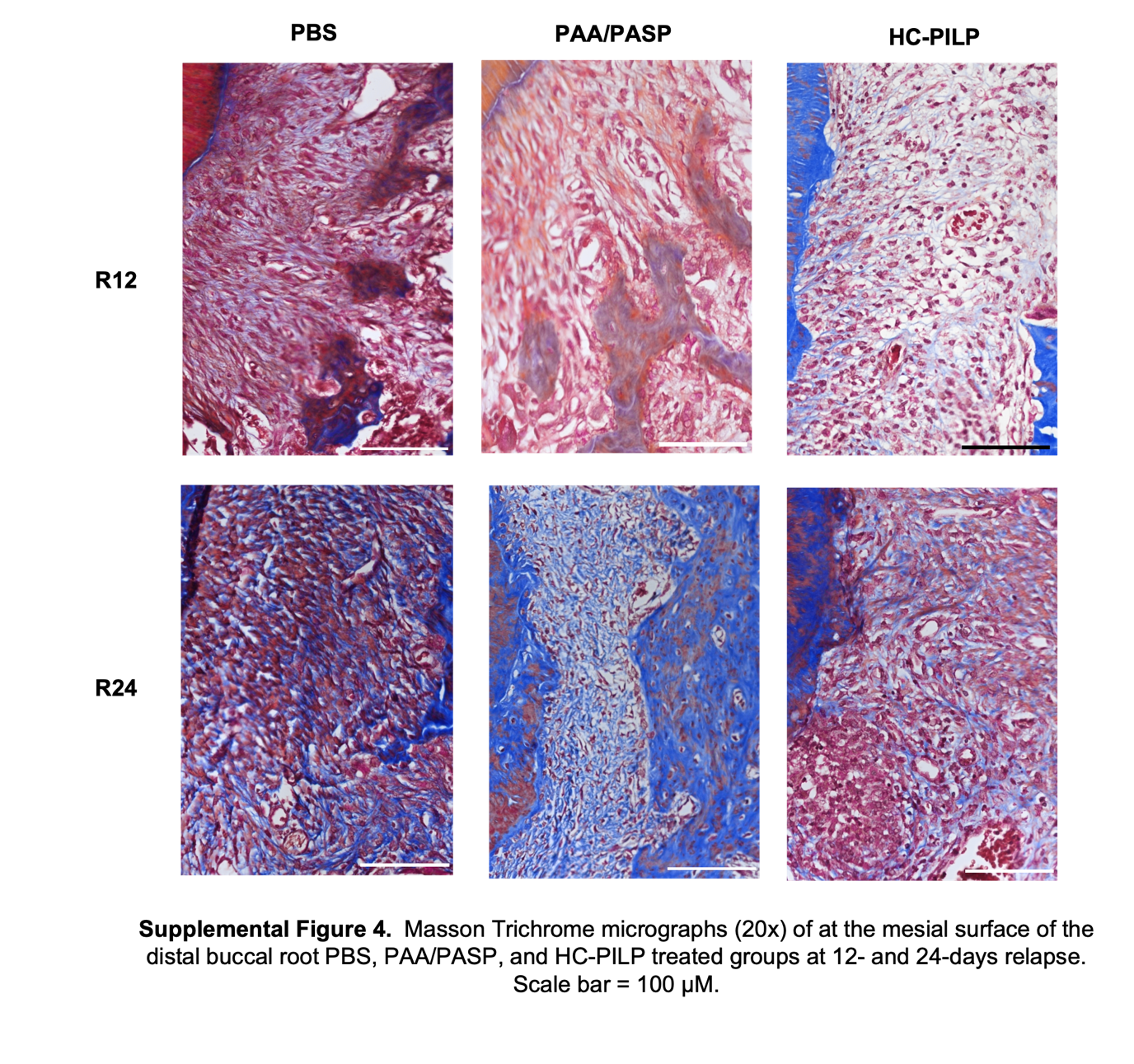

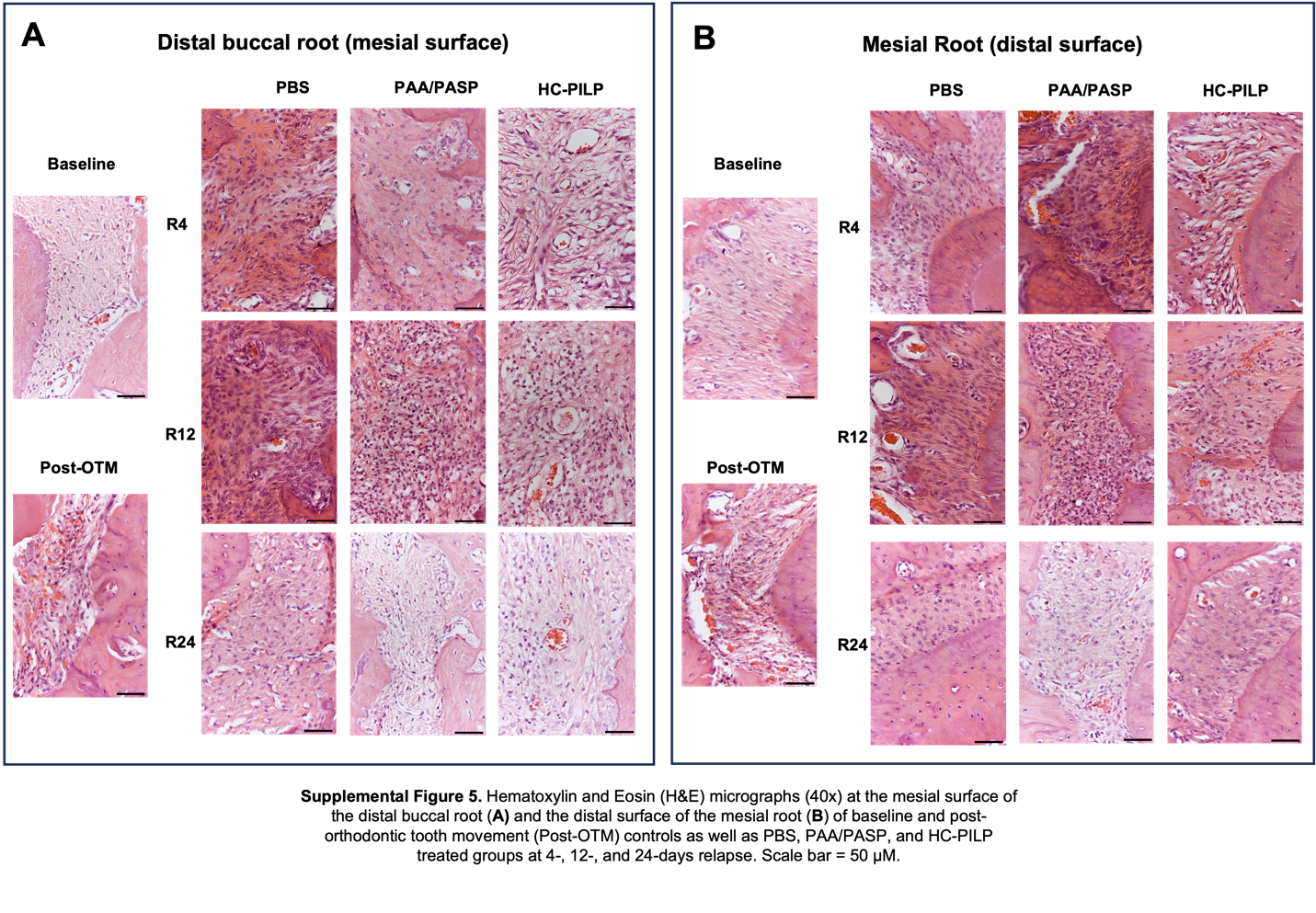

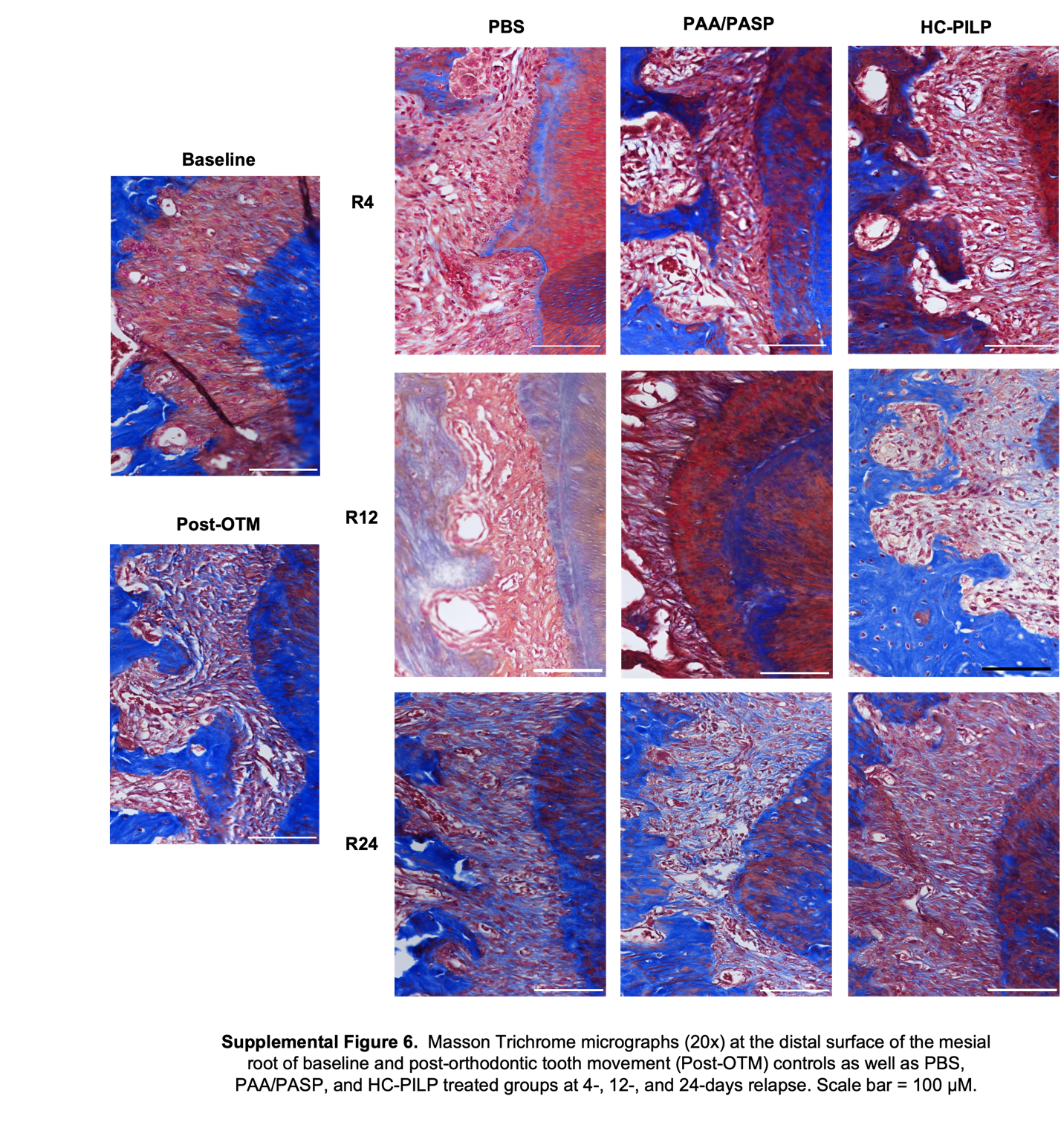

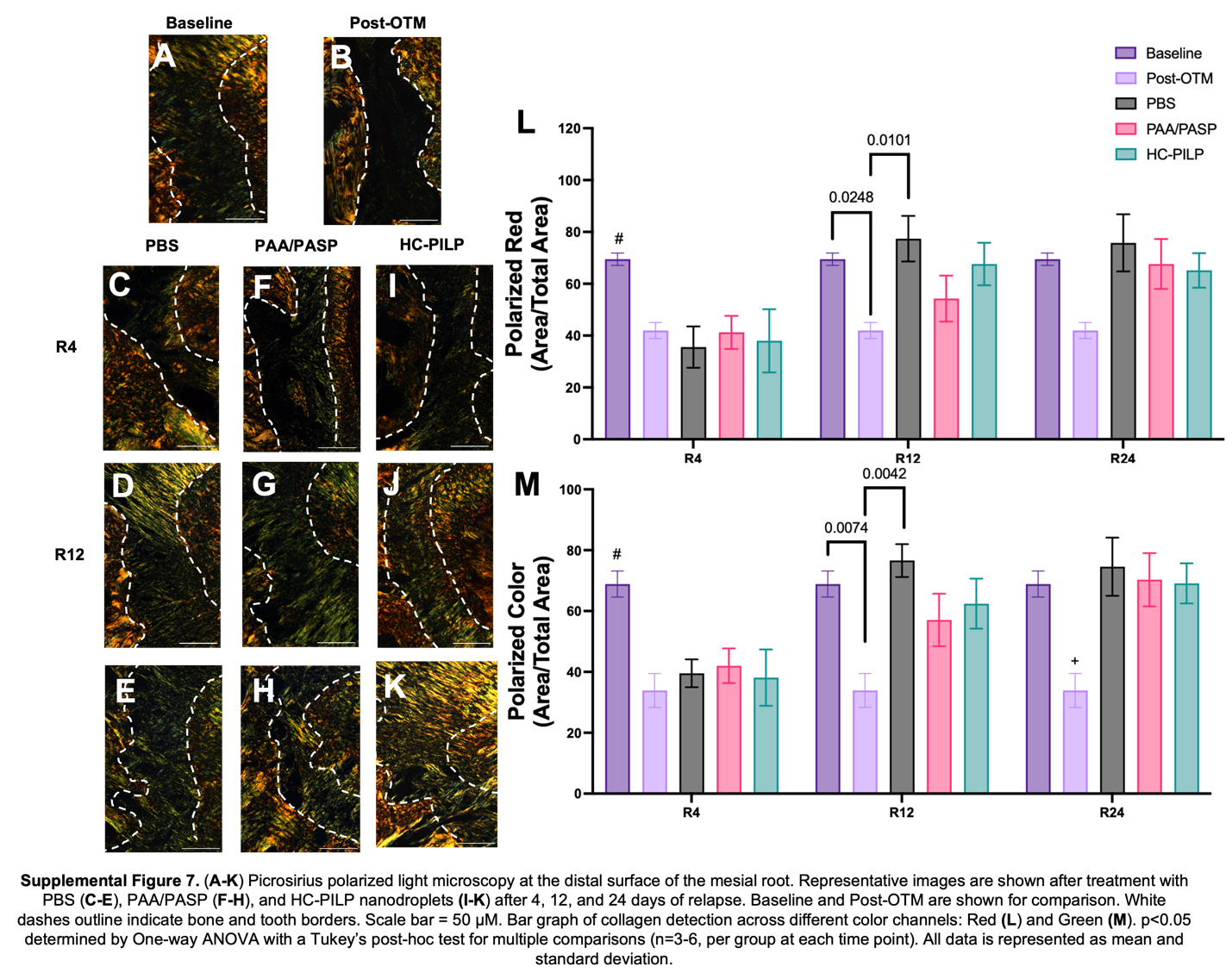

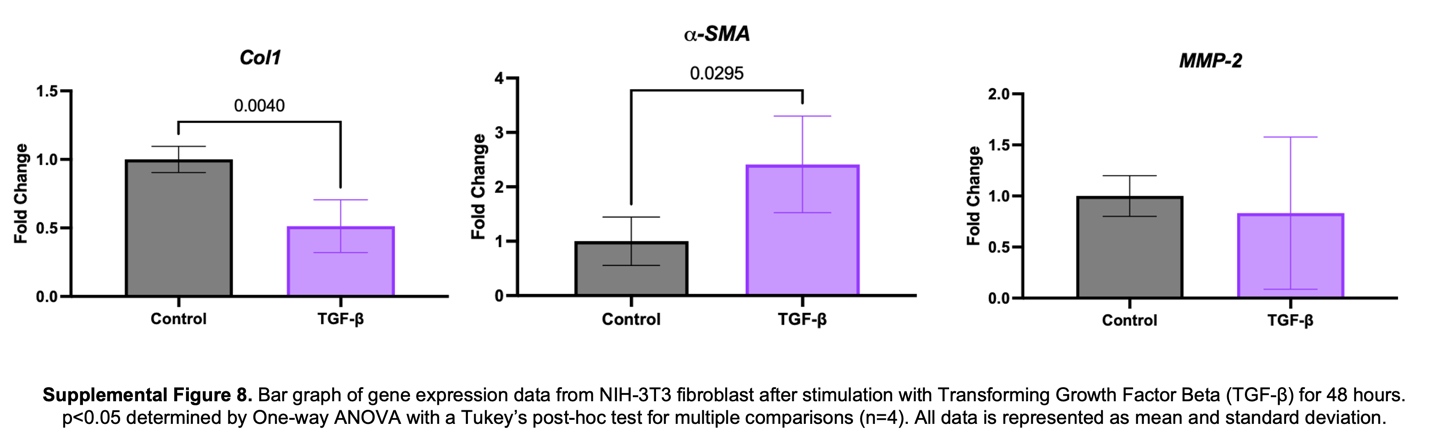

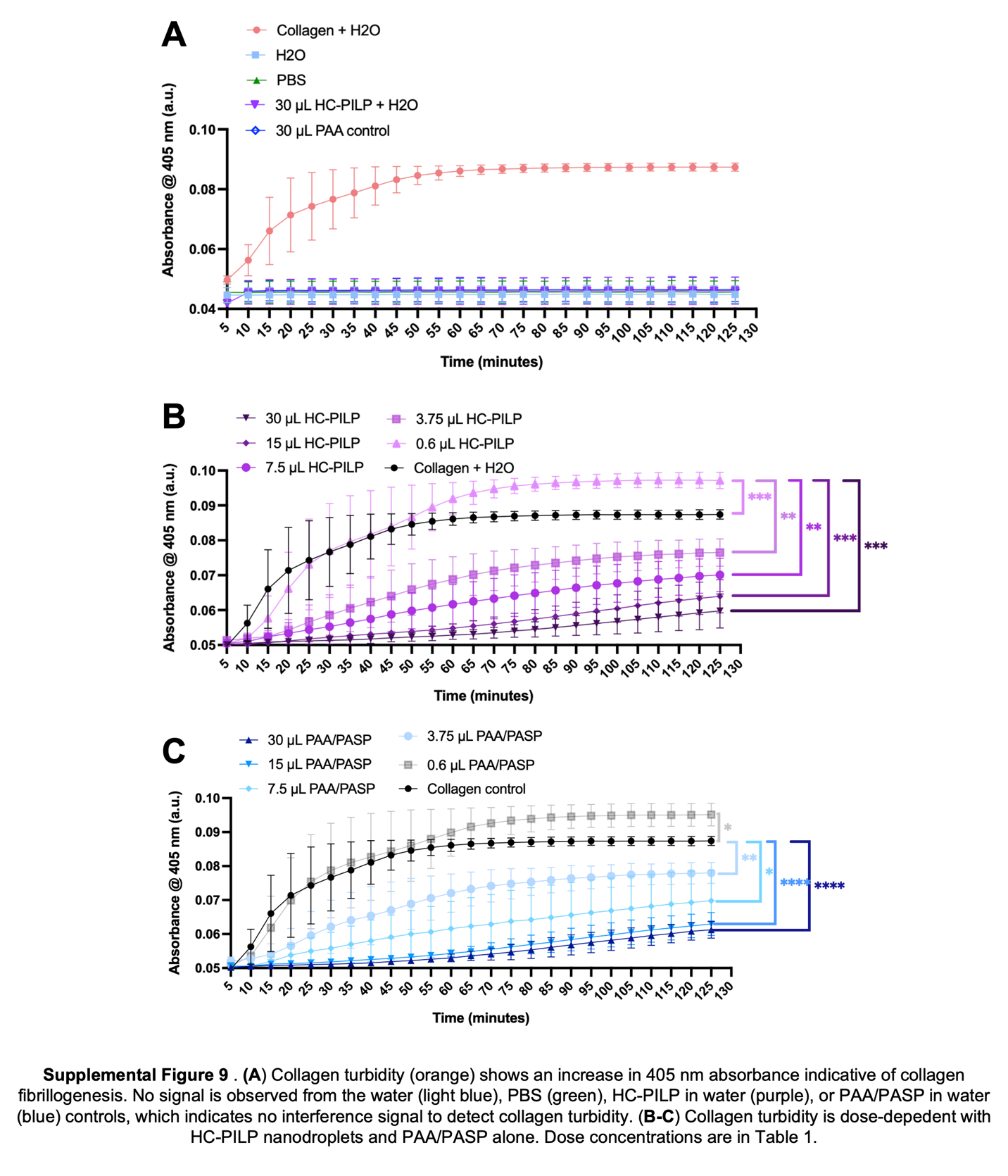
